# Rapidly Increasing SARS-CoV-2 Neutralization by Intravenous Immunoglobulins Produced from Plasma Collected During the 2020 Pandemic

**DOI:** 10.1101/2021.02.12.430933

**Authors:** Maria R. Farcet, Michael Karbiener, Julia Schwaiger, Reinhard Ilk, Thomas R. Kreil

## Abstract

Immunoglobulin (IG) lots (N=176) released since March 2020 were tested for SARS-CoV-2 neutralizing antibodies, with first positive results for September 2020 lots, mean = 1.8 IU/ml, 46% of lots positive. From there, values steadily increased, in correlation with the cumulative COVID-19 incidence, to reach a mean of 36.7 IU/ml and 93% of lots positive by January 2021. Extrapolating the correlation, IGs could reach an anti-SARS-CoV-2 potency of ~400 IU/ml by July 2021. At that stage, prophylactic IG treatment for primary/secondary immunodeficiency could contain similar doses of anti-SARS-CoV-2 as convalescent plasma which is used for treatment of COVID-19.

## Background

People with primary and secondary immunodeficiencies (PID / SID) need substitution therapy with antibodies prepared from the plasma of healthy donors in the form of immunoglobulin (IG) preparations. For emerging agents, however, the plasma donor community is initially seronaïve, and thus IG preparations cannot afford protection against these new infectious agents.

After the emergence of another zoonotic Coronavirus in humans, Severe acute respiratory syndrome coronavirus-2 (SARS-CoV-2), it was initially unclear whether past infection with other, seasonally circulating Human coronaviruses (HCoV) would have induced cross-protective antibodies to the new virus. Using the gold standard virus neutralization test this was shown not to be the case, i.e. intravenous immunoglobulins (IVIG) produced from plasma collected before the Corona virus disease 2019 (COVID-19) pandemic did not neutralize SARS-CoV-2 [1].

By the end of 2020, close to 100 million COVID-19 cases had been reported globally, with almost a quarter of them in the US alone [2]. The US is quantitatively the most important origin of plasma for fractionation [3], and thus an increasing proportion of the plasma supply will be collected from donors post COVID-19, plasma that is now expected to also carry antibodies to SARS-CoV-2.

Here, we report the results from an investigation into the seroconversion of the US supply of plasma for fractionation, through testing of IVIGs derived exclusively from US plasma for SARS-CoV-2 neutralization, which revealed rapidly increasing antibody titers which correlated with cumulative COVID-19 incidence. Together with the characterization of SARS-CoV-2 neutralizing antibody (nAb) titers in a large collection of COVID-19 convalescent plasma (CP) donations, this permits a near-term projection of potentially protective SARS-CoV-2 antibody levels in future IG lots, particularly when used in a prophylactic setting for substitution therapy.

## Methods

### Immunoglobulin preparations

A total of 176 IVIG lots (Gammagard Liquid; Baxter Healthcare Corp., Westlake Village, CA) released between March 2020 and January 2021 from plasma collected by plasmapheresis (source plasma) in the US were analyzed. For a subset of 12 of these IVIG lots, information about the dates of plasma collection was obtained.

### COVID-19 convalescent plasma samples

438 COVID-19 CP samples collected between March and July 2020 were obtained from Austrian (n = 300) and US (n = 138) plasma donation centers (BioLife). The samples originated from donors who had PCR-confirmed SARS-CoV-2 infections or who described a disease progression consistent with COVID-19 and for which SARS-CoV-2 neutralization was confirmed. Information on disease severity was requested from each donor. Donors signed informed consent and agreed to additional testing.

### Measurement of SARS-CoV-2 and HCoV-229E neutralizing antibodies

SARS-CoV-2 and HCoV-229E neutralizing antibody (nAb) titers were determined using materials and methods previously reported [1]. Briefly, 2-fold serially diluted samples were incubated with equal volumes of SARS-CoV-2 (strain “BavPat1/2020”, Charité Berlin, Germany) or HCoV-229E (for IVIG samples, Cat. no. VR-740, ATCC, Rockville, MD) at 10^3.0^ tissue culture infectious doses 50% per milliliter (TCID_50_/ml) and incubated for 150 min before titration on Vero cells (for SARS-CoV-2; Cat. no. 84113001, ECACC, Porton Down, Salisbury, UK) or MRC-5 cells (for HCoV-229E; Cat. no. CCL-171, ATCC) in eight-fold replicates per dilution. The virus-induced cytopathic effect was determined after 5-7 days of incubation. The reciprocal sample dilution resulting in 50% virus neutralization (NT_50_; SARS-CoV-2 detection limit in IVIG: ≤ 1:2) was determined using the Spearman-Kärber formula, and the calculated neutralization titer for 50% of the wells reported as 1:X. For further analyses, samples with a neutralization titer below the detection limit were assigned a value of 0.5x the detection limit. The National Institute of Biological Standards and Control (NIBSC, Potters Bar, UK) research reagent 20/130, for which a potency in international units has recently been assigned [4], was included in the study and the concentration of SARS-CoV-2 nAbs therefore reported in IU/ml.

Testing was done using a fully validated analytical method (for SARS-CoV-2 nAbs) or a controlled assay that included several validity criteria, i.e. confirmatory titration of input virus infectivity and cell viability (for HCoV-229E nAbs).

### Graphs and statistical analysis

Overall COVID-19 incidence in the US was taken from the Centers for Disease Control and Prevention (CDC) COVID data tracker [2]. Data analysis and visualization was done using GraphPad Prism v8.1.1 (San Diego, CA), R Studio v1.1.383 (Boston, MA), Minitab v. 17.3.1 (State College, PA) and Microsoft Excel.

Mean SARS-CoV-2 antibody values for IVIG lots released in a month were correlated against the cumulative incidence of COVID-19 in the US that was recorded 6 months prior to IVIG release, e.g. the mean antibody potency measured for IVIG lots released in September 2020 was correlated against cumulative COVID-19 incidence in the US in March 2020. A log-transformed linear model and a polynomial regression model indicated comparable quality of fit and both were used to calculate an extrapolation beyond the period for which antibody measurements were made. The slope at the highest observed cumulative incidence (i.e. July 2020; 1.39%) was used for extending the models in a linear manner.

## Results

### SARS-CoV-2 and HCoV-229E neutralizing antibodies in commercial IVIG preparations

SARS-CoV-2 nAbs were undetectable for IVIG lots released to the market between March and August 2020 (n = 63). For IVIG lots released in September 2020, 12 of 26 lots (46%) were seropositive with a mean SARS-CoV-2 nAb concentration of 1.8 IU/ml (Figure 1A). From there onwards, the proportion of SARS-CoV-2 nAb positive IVIG lots steadily increased, with mean nAb concentrations of 3.0 (n = 7/13; 54%) in October, 4.8 (n = 20/30; 67%) in November, 12.1 (n = 16/17; 94%) in December, and 36.7 (n = 25/27; 93%) IU/ml for January 2021, respectively (Figure 1A).

**Figure 1.**
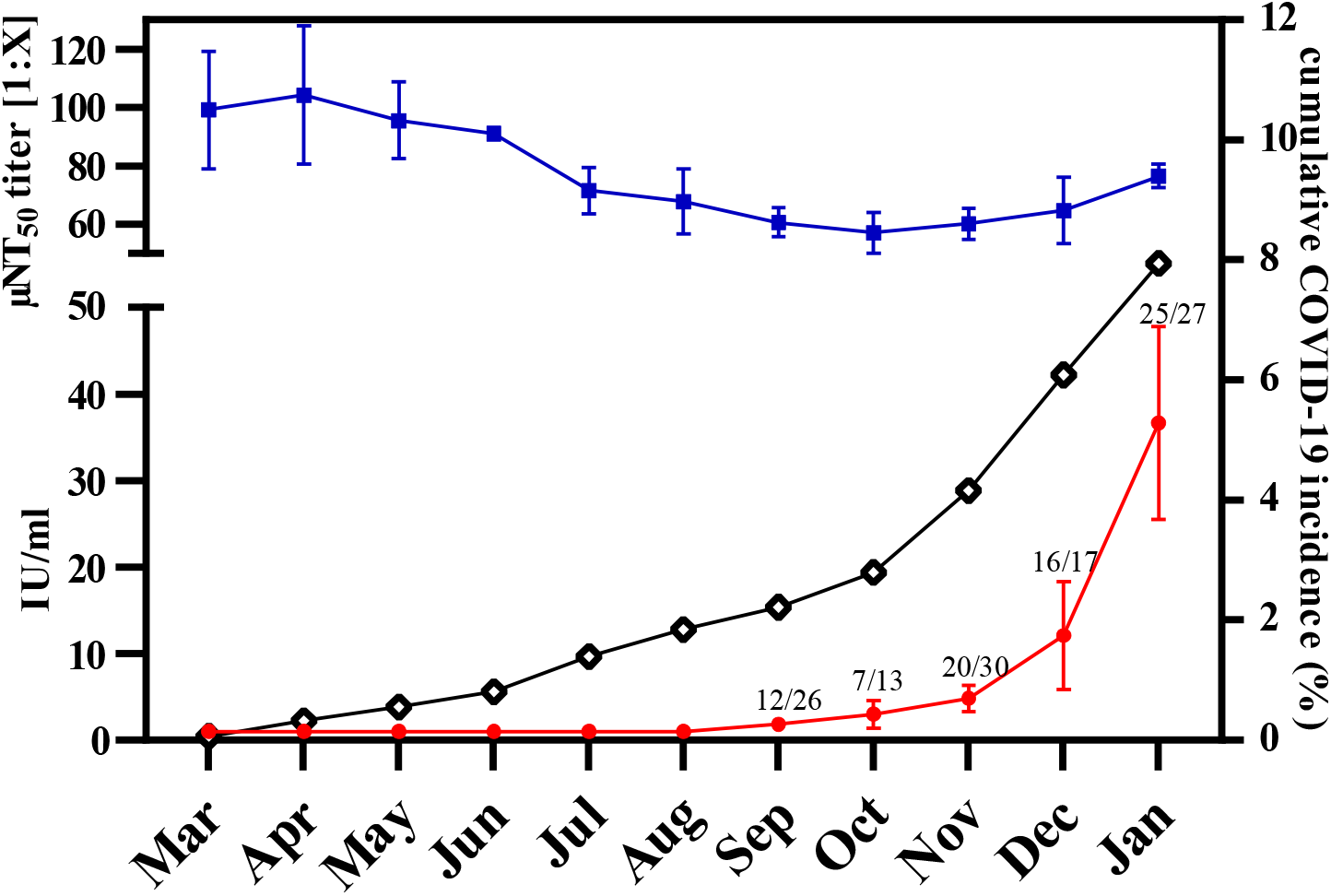
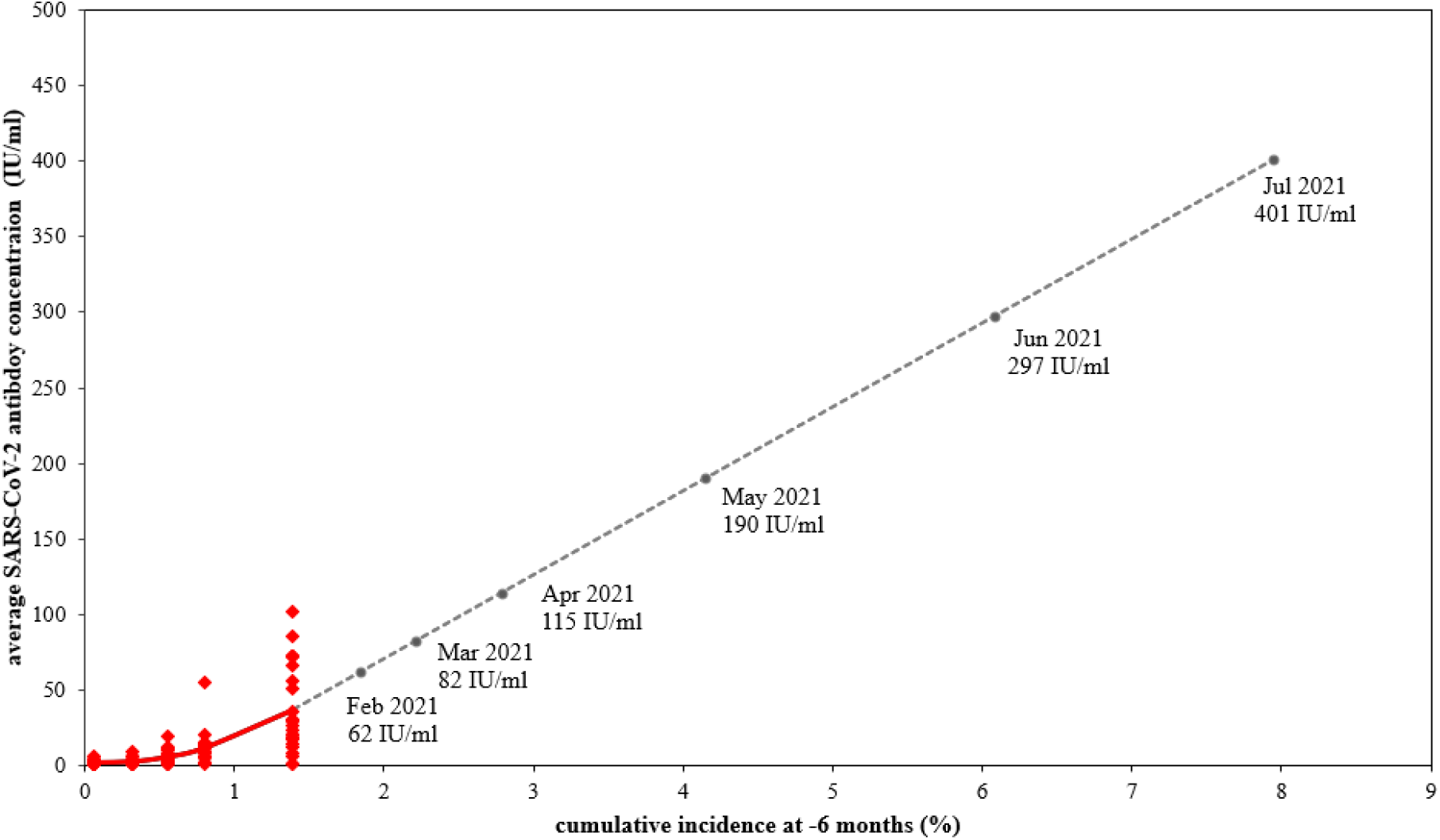
**(A)** SARS-CoV-2 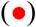 neutralizing antibody concentration and Human coronavirus 229E 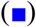 neutralizing antibody titers in 176 commercial IVIG lots manufactured from March 2020 until January 2021; mean values ± 95% confidence interval are indicated, the number of IVIG lots for which SARS-CoV-2 neutralization was detected and the total number of IVIG lots tested are indicated for each month. SARS-CoV-2 incidence (◊) in the US population (confirmed cases per million) is shown. **(B)** Prediction model of SARS-CoV-2 antibody development in future IVIG lots. Significant curvilinearity (red sold line; P-value = 0.006) was seen in the relationship between −6 months COVID-19 cumulative incidence in the US population and measured SARS-CoV-2 antibody concentrations in IVIG lots 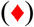. The slope at the upper end of the observation range was used for linear extrapolation (grey dotted line) of SARS-CoV-2 antibody concentrations in IVIG lots that will be released in the coming months 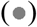. Abbreviations: IVIG, intravenous immunoglobulin; IU/ml, international units per milliliter; μNT50, 50% neutralization titer; SARS-CoV-2, Severe acute respiratory syndrome coronavirus 2.

HCoV-229E nAb titers of the same IVIG lots remained at similar levels throughout the period surveyed (Figure 1A) and the observed variations in HCoV-229E nAb titers were similar to previously observed ranges [1].

### COVID-19 incidence in the US and development of SARS-CoV-2 antibody content

For the first 12 IVIG lots that contained measurable SARS-CoV-2 nAbs, the time of collection for the several thousand plasma units used for production was analyzed. On average, plasma was collected six months before IVIG lot release. Thus, the cumulative incidence as the underlying cause for the seroprevalence in any given month would be expected to determine the level of antibodies present in IVIG lots released six months later. The low SARS-CoV-2 antibody content of IVIG lots released in September 2020 would thus on average be derived from plasma collected in March 2020, when the cumulative COVID-19 incidence was 0.06% of the US population (Figure 1A). By July 2020, the cumulative incidence had increased to 1.39%, i.e. about 20-fold, and similarly the mean SARS-CoV-2 antibody concentration increased from 1.8 IU/ml for September 2020 IVIG lots, to 36.7 IU/ml for IVIG lots released in January 2021 (Figure 1A). Based on the increasing cumulative COVID-19 incidence since August 2020, an estimation of SARS-CoV-2 nAb concentrations in future IVIG lots is possible (Figure 1B). Given a cumulative incidence of 7.94% in January 2021, IG lots to be released in July 2021 are expected to contain a mean SARS-CoV-2 nAb concentration of around 400 IU/ml (Figure 1B).

### SARS-CoV-2 neutralizing antibodies in COVID-19 convalescent plasma samples

A large collective of COVID-19 CP units (n = 438) was tested with the gold standard neutralization assay for anti-SARS-CoV-2 potency and the results reported in relation to the newly assigned WHO standard [4]. The mean SARS-CoV-2 nAb concentration was 301 IU/ml (Figure 2: IU/ml histogram, median 156, 20^th^ percentile 75, 80^th^ percentile 353, range < 2 - 6,937). A mean of 516 IU/ml was determined for all units above the median (n = 218) and a mean of 945 IU/ml for the top 20% of CP characterized in this study (n = 88). No SARS-CoV-2 nAbs were detected in the sample of one donor who had PCR-confirmed SARS-CoV-2 infection. Most donors had experienced asymptomatic or mild COVID-19 (75% of samples), around 9% of samples originated from donors who had recovered from severe COVID-19.

**Figure 2.**
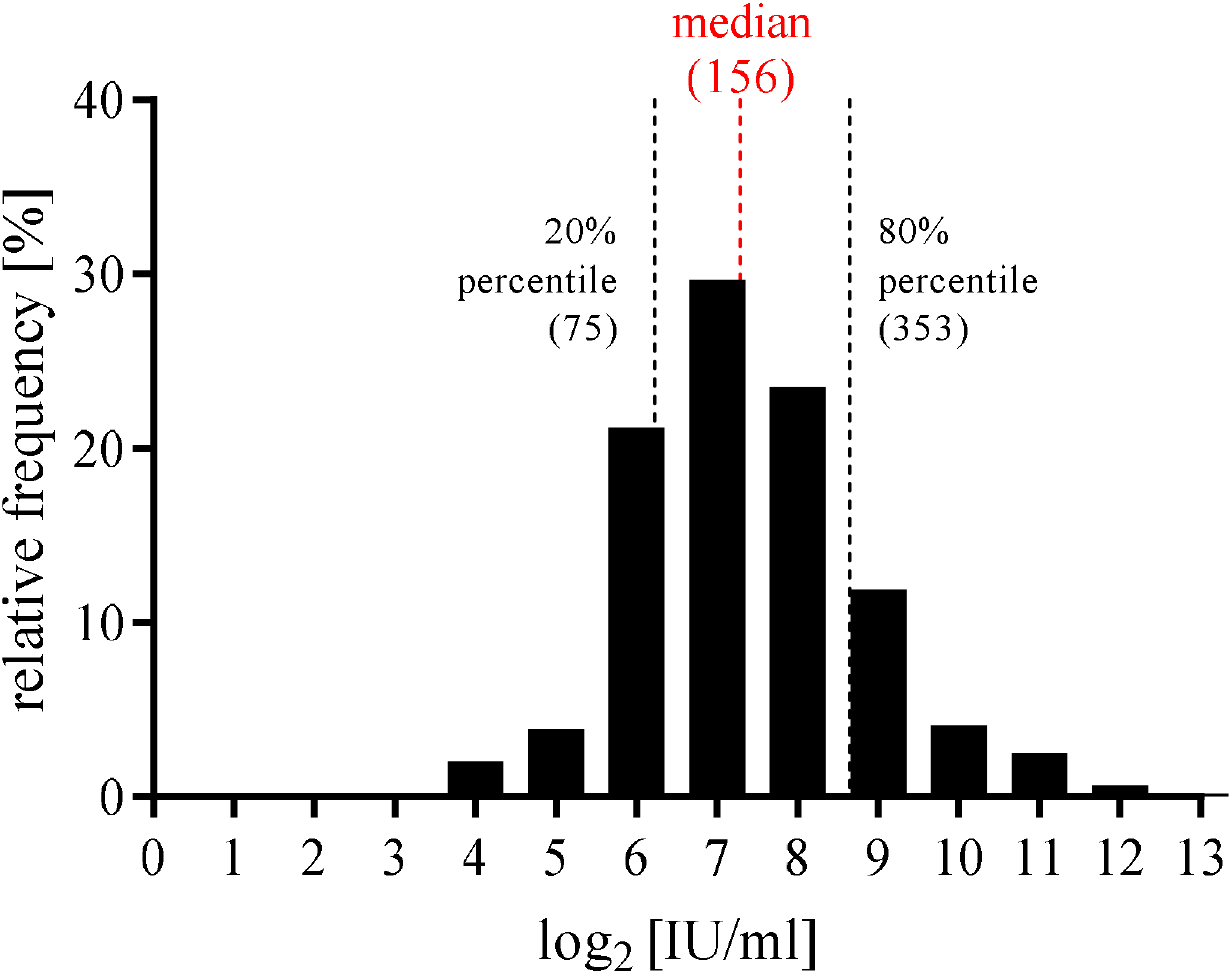
Characterization of 438 COVID-19 convalescent plasma samples collected in Austrian and US BioLife Plasma centers. SARS-CoV-2 neutralizing antibody content is reported as log_2_ [IU/ml] against relative frequency of occurrence (%). Median, 20% and 80% percentile values are indicated as IU/ml. Abbreviations: IU/ml, international units per milliliter; SARS-CoV-2, Severe acute respiratory syndrome coronavirus 2.

## Discussion

The cumulative incidence of past COVID-19 cases, i.e. the proportion of individuals expected to be SARS-CoV-2 antibody positive, in the US in any given month forms the basis for seropositivity in IG lots released to the market approximately six months later. This prediction assumes a constant proportion of asymptomatic infections to COVID-19 cases, as they, too, are expected to result in antibody positive plasma donors.

For the extrapolation of future SARS-CoV-2 neutralizing antibody concentrations in IG, consideration of only past COVID-19 cases is very conservative. With large vaccination campaigns under way at this moment, approximately 9.6% of the US population has already been vaccinated by the end of January 2021 [2]. Vaccine-induced antibodies will thus further increase the anti-SARS-CoV-2 potency of future lots of IGs, and based on the current cumulative incidence of 7.94% in addition to a 9.6% vaccination rate, and higher mRNA vaccine-induced antibody titers than post-COVID [5, 6], the anti-SARS-CoV-2 potency of future IG products may potentially be higher than the SARS-CoV-2 potency extrapolated here (Figure 1B).

To date, the level of SARS-CoV-2 neutralizing antibody titers required in IG to provide protection against COVID-19 has not been determined. A comparison of the antibody potency contained in CP, to those expected in IG soon may provide for some perspective.

In a large cohort of COVID-19 convalescents, a median anti-SARS-CoV-2 potency of 156 IU/ml was determined (Figure 2). The original US FDA emergency use authorization for transfusion of CP required use of plasma at a potency of above the median, as determined by a high throughput binding assay [7, 8]. The mean neutralization potency of above median units of the cohort of samples tested here is 516 IU/ml. Joyner et al. [7] did, however, report significantly better clinical success when using CP above the 80^th^ percentile, the mean of which in neutralization potency is 945 IU/ml. With about 200 ml used for CP transfusion, this would equate to SARS-CoV-2-neutralizing antibody doses of 103,200 IU and 189,000 IU, respectively.

By July 2021, IGs can be extrapolated to contain a mean potency of approximately 400 IU/ml, of which standard prophylaxis regimen for PID / SID would apply approximately 500 mg IG/kg, resulting in the administration of 350 ml IG for a 70 kg person, or a dose of 140,000 IU. While the total doses would be quite similar, it has become evident that CP treatment was more successful when administered at early stages of COVID-19 [9, 10], i.e. before extensive virus spread within respiratory and other organs. Regular IG substitution therapy for the treatment of PID and SID represents prophylaxis, i.e. antibody administration even before virus exposure, and thus should have a significantly better likelihood of success. This prospect is of particular importance for PID patients, for whom a 10-fold higher COVID-19 mortality rate has been reported [11]. The above calculations are quite conservative, as in the US already now the proportion of vaccinated individuals has surpassed those with post COVID-19, a trend that can be expected to accelerate in the months to come. In addition, currently available data indicate superior levels of SARS-CoV-2 antibodies after mRNA-based vaccines as currently in use in the US [5, 6]. Cumulatively, these circumstances would appear to support rapidly increasing levels of SARS-CoV-2 antibodies in IG, so that IG-mediated protection against COVID-19 for regularly substituted PID / SID seems quite possible. More research to confirm the extrapolations from this study is currently under way. For COVID-19 treatment, rather than prophylaxis, a hyper-IVIG manufactured exclusively from plasma of COVID-19 convalescent donors is currently under evaluation in a phase III clinical trial [12].

## Acknowledgments

The contributions of the entire Global Pathogen Safety team, most notably Melanie Graf, Simone Knotzer, Jasmin de Silva, Stefan Pantic, Brigitte Kainz, Julius Segui (neutralization assays), Veronika Sulzer, Sabrina Brandtner (cell culture), Eva Ha and Alexandra Schlapschy-Danzinger (virus culture), Kathrin Nagl (IVIG logistics) as well as Eva Gschaider-Reichhart (data collection) are gratefully acknowledged. Harold Batista (production planning) compiled plasma donation dates. Colleagues from BioLife Austria and US, especially Nadja Frankl and Kathryn Barrett supplied CP samples. SARS-CoV-2 was sourced via EVAg (supported by the European Community) and kindly provided by Christian Drosten and Victor Corman (Charité Universitätsmedizin, Institute of Virology, Berlin, Germany).

